# Bioinformatic analysis of placental exomiRs targeting the brain in preeclampsia

**DOI:** 10.1101/2025.10.06.680827

**Authors:** Marcelo Alarcón, Claudio Aguayo, Manu Vatish, Carlos Escudero

**Author notes:** Correspondence: Marcelo Alarcón Thrombosis and Healthy Aging Research Center Group of Research and Innovation in Vascular Health Medical Technology School Department of Clinical Biochemistry and Immunohematology Faculty of Health Sciences Universidad de Talca Talca, Chile. Carlos Escudero, MD PhD Vascular Physiology Laboratory Group of Research and Innovation in Vascular Health Basic Sciences Department Faculty of Sciences Universidad del Bio-Bio Chillán, Chile /.

## Abstract

**Background:** Preeclampsia is a hypertensive disorder of pregnancy associated with systemic endothelial dysfunction and, in severe cases, maternal neurological complications. Placenta-derived exosomal microRNAs (exomiRs) mediate inter-organ communication and may contribute to neurovascular injury, but their role in maternal brain dysregulation remains unclear.

**Methods:** We performed systems-level bioinformatic analyses of 12 differentially expressed exomiRs (6 from early-onset and 6 from late-onset preeclampsia) to assess their potential impact on brain homeostasis. Target interactomes were examined for functional enrichment, subcellular localization, and network complexity. ExomiR targets were integrated with cerebrospinal fluid (CSF) proteomic profiles from preeclamptic women with neurological symptoms. We further evaluated regulation of blood–brain barrier (BBB) components and mapped spatial expression of target proteins across brain regions and cell types relevant to neurovascular function.

**Results:** Early-and late-onset preeclampsia exomiRs displayed distinct interactomes and biological signatures. Early-onset exomiRs were linked to RNA metabolism, oxidative stress, and endothelial stability, whereas late-onset exomiRs were enriched in nitrogen metabolism and vesicle-mediated transport. Integration with CSF proteomics revealed convergence on dysregulated proteins, including MAPK8, RTN4, and YWHA family members, involved in inflammation, neurodegeneration, and axonal inhibition. Several exomiRs targeted BBB-related proteins (CLDN2, TJP1, EFNA5), suggesting coordinated barrier disruption. Spatial mapping localized these targets to endothelial cells, astrocytes, interneurons, and pyramidal neurons, implicating altered synaptic and glial regulation.

**Conclusions:** Preeclampsia-associated exomiRs may impair maternal brain homeostasis through coordinated regulation of neurovascular, inflammatory, and synaptic pathways. These findings identify candidate molecular mediators of preeclampsia-related neurovascular dysfunction and potential biomarkers for maternal brain injury.

## Introduction

Preeclampsia (PE) is a multifactorial hypertensive disorder of pregnancy that typically arises after the 20th week of gestation. It is defined by new-onset hypertension accompanied by proteinuria or signs of maternal organ dysfunction and affects approximately 2–8% of pregnancies globally (Giannubilo et al., 2024). Despite improvements in prenatal care, PE remains a leading cause of maternal and fetal morbidity and mortality (Shan et al., 2024).

PE has diverse clinical manifestations, including severe cerebrovascular complications. Eclampsia, the most dramatic, is characterized by the new onset of generalized tonic–clonic seizures in pregnancy, typically in the context of hypertension and proteinuria. Other neurological complications include ischemic and hemorrhagic stroke, as well as cerebral edema, all associated with high maternal mortality (Escudero et al., 2023; Younes and Ryan, 2019). Moreover, women with a history of PE face a significantly increased risk of developing long-term neurological conditions, particularly vascular dementia and Alzheimer’s disease (Carey et al., 2024; Schliep et al., 2023).

Increasing evidence highlights the role of extracellular vesicles (EVs), especially exosomes, in mediating maternal-placental communication during pregnancy (Ghosh et al., 2024). Exosomes are nano-sized (30-150 nm) lipid-bound vesicles actively released by placental trophoblasts into the maternal circulation. These vesicles carry a variety of molecular cargo, including proteins, lipids, mRNAs, and microRNAs (miRNAs) (Ghosh et al., 2024). MiRNAs are ∼22-nucleotide-long non-coding RNAs that regulate gene expression post-transcriptionally, primarily by targeting complementary sequences on messenger RNAs and inhibiting their translation or promoting degradation (Pillay et al., 2017).

In PE pregnancies, the composition of placental exosomes is significantly altered. Several miRNAs are differentially expressed in exosomes derived from PE placentas, such as miR-210-3p, miR-193b-5p, and miR-9-5p. These miRNAs are implicated in pathways affecting angiogenesis, mitochondrial function, inflammation, and trophoblast invasion (Gao et al., 2020). For example, miR-210-3p is hypoxia-inducible and suppresses mitochondrial respiration while promoting oxidative stress, both of which are central elements of PE pathophysiology (Awoyemi et al., 2024). Beyond their role in placental function, PE-associated miRNAs also target genes involved in key neurological functions, including synaptic plasticity, neurogenesis, and mitochondrial metabolism. Therefore, dysregulation of these pathways may help explain observations of acute and long-term consequences observed in affected women (Suvakov et al., 2023).

Importantly, exosomal miRNAs (exomiRs) can cross biological barriers, including the blood-brain barrier (BBB), and exert effects in distant organs such as the brain (Alvarez-Erviti et al., 2011). Animal and *in vitro* studies demonstrate that EVs derived from preeclamptic placentas disrupt the BBB integrity by downregulating tight junction proteins (e.g., claudin-5), thereby increasing endothelial permeability (Sandoval et al., 2025). This increased permeability likely promotes persistent vascular dysfunction, neuroinflammation, and the transfer of neurotoxic molecules—including misfolded proteins and EV-associated miRNAs—similar to mechanisms described in chronic neurological diseases such as Alzheimer’s disease (Xu et al., 2021).

Some PE-associated miRNAs, such as miR-93-5p, miR-31-5p, and miR-92a-3p, can be detected as early as the first trimester, offering potential for early identification of high-risk pregnancies (Awoyemi et al., 2024). These miRNAs not only reflect underlying placental pathology but also participate in the dysregulation of angiogenic signaling and inflammatory pathways (Shan et al., 2024). Systems biology analyses further reveal that PE-related miRNAs converge on PI3K/AKT, VEGF, and NF-κB pathways, thereby linking placental dysfunction to systemic endothelial activation (Ghosh et al., 2024) and potentially to cerebral endothelial injury.

Proteomic analyses of cerebrospinal fluid (CSF) from women with PE have revealed differentially expressed proteins that converge on vascular and inflammatory pathways (Ciampa et al., 2018; van den Berg et al., 2017). However, no direct associations have yet been established between placental EV– derived miRNAs and brain-related protein targets. This gap suggests that circulating placental exomiRs could represent valuable biomarkers for systemic maternal alterations, particularly cerebrovascular complications, in women affected by PE.

This study bioinformatically explored how placenta-derived exomiRs may contribute to maternal brain dysregulation. Also, we integrated exomiR target networks with cerebrospinal fluid proteomics and brain region–specific protein expression to identify candidate molecular mediators of neurovascular dysfunction in preeclampsia.

## Materials and Methods

### ExomiR selections

To investigate the pathophysiological role of exomiRs in PE, we analyzed exomiRs reported by Pillay et al. (2019) as differentially expressed in early-onset PE (EOPE, <34 weeks of gestation) and late-onset PE (LOPE, ≥34 weeks), compared with gestationally matched normotensive controls (Pillay et al., 2019).

### Candidate targets collection and Protein-Protein Interactions (PPIs) Network Construction

Predicted and validated targets of selected exomiRs were retrieved from multiple databases, including miRBase (Kozomara et al., 2019), miRDB (Liu and Wang, 2019), and Target Scan Human (Agarwal et al., 2015). Protein–protein interactions (PPIs) and tissue expression patterns were obtained from the Integrated Interactions Database (IID) (Pastrello et al., 2020) and the Human Protein Atlas (Sjostedt et al., 2020), respectively. The targets associated with PE were determined using an online Venn diagram tool to visualize the overlapping results. In the constructed interactome, first-order proteins were defined as direct exomiRs targets, while second-order proteins were defined as those regulated by the first-order proteins (Figure 1A).

**Figure 1.**
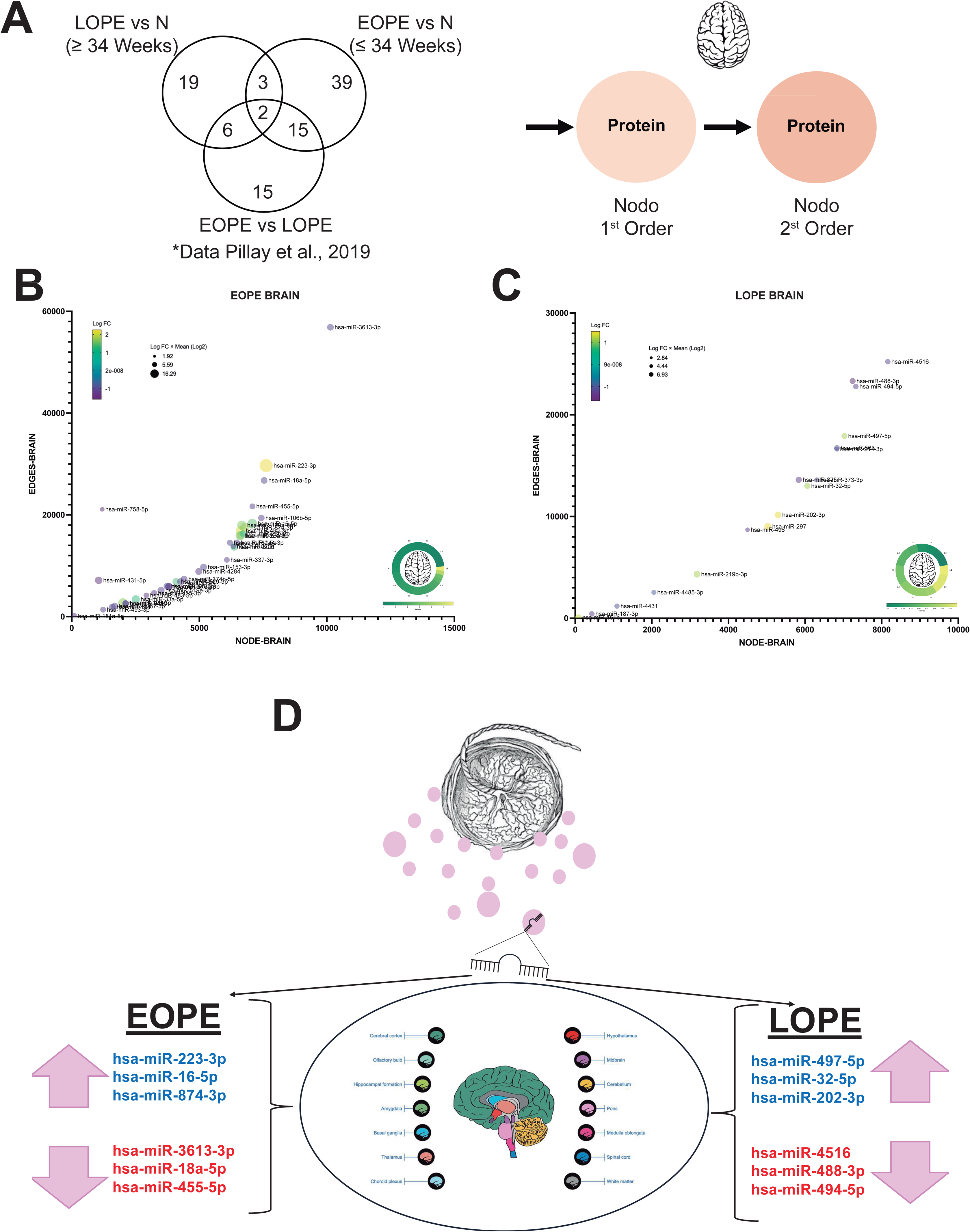
Brain-specific network architecture of EOPE and LOPE-associated exomiRs. (A) Venn diagram of differentially expressed placental exomiRs among EOPE vs. controls (≤33 weeks), LOPE vs. controls (≥34 weeks), and EOPE vs. LOPE comparisons (adapted from Pillay et al., 2019). (B) Each dot represents an EOPE-associated exomiR; size reflects network node count, and color intensity indicates the “Log FC × Mean (Log₂)” value. (C) Each symbol corresponds to a LOPE-associated exomiR; the position reflects the network size and the “Log FC × Mean (Log₂)” value. (D) Selection of 12 brain-relevant exomiRs (six EOPE, six LOPE) based on network complexity and differential expression, with up- and down-regulated miRNAs positioned to emphasize their distinct contributions to early-versus late-onset PE cerebral pathology.

To further analyze the results from these databases, Cytoscape 3.7.2 was used as a tool to visualize and analyze the PPI network (Shannon et al., 2003).

### Enrichment Biological function

The Gene Ontology (GO) analysis enrichment was performed using ShinyGO 0.81 database (Ge et al., 2020). This enabled systematic characterization of exomiR-regulated targets across biological processes (BP), molecular functions (MF), and cellular components (CC), as well as the identification of relevant signaling pathways and regulatory mechanisms.

### Brain-Specific Protein Mapping

To assess the potential impact of selected exomiRs on the central nervous system, we conducted a spatial mapping analysis of exomiR-regulated targets across human brain regions. Twelve exomiRs, chosen based on differential expression and network complexity, were analyzed against proteomic data from the Human Protein Atlas (HPA) (Thul and Lindskog, 2018). Protein expression was examined 14 distinct anatomical regions of the brain: amygdala, basal ganglia, cerebellum, cerebral cortex, choroid plexus, hippocampus, hypothalamus, medulla oblongata, midbrain, olfactory bulb, optic tract, pons, spinal cord, thalamus, and white matter. Predicted exomiR targets were cross-referenced with the HPA dataset to identify regionally expressed proteins under putative post-transcriptional regulation.

### Subcellular Localization of ExomiR-Regulated Proteins

To determine the specific protein targets regulated by each exomiR and to characterize their subcellular distribution, we performed a multi-database annotation. First, experimentally validated or high-confidence predicted targets for each exomiR were compiled (see Candidate targets section). We then queried three curated databases to retrieve subcellular localization information for each protein: the Human Protein Atlas (Thul and Lindskog, 2018), UniProt (Ahmad et al., 2025), and COMPARTMENTS database (Binder et al., 2014). Only proteins with reported or strongly predicted subcellular localizations in at least one of the databases were retained for downstream analyses. This approach enabled us to classify exomiR targets according to their presence in major cellular compartments, including the centrosome, cytoskeleton, cytosol, endoplasmic reticulum, endosomes, Golgi apparatus, lysosomes, mitochondria, nucleus, peroxisomes, plasma membrane, secreted, and vesicular compartments. Proteins with no defined localization in any of the three databases were categorized as “unmapped”. This standardized classification enabled a comprehensive assessment of the intracellular distribution of exomiR-regulated proteins, providing insight into their potential mechanistic roles in post-transcriptional and inter-organelle regulation.

## Results

### Systemic and Brain-Specific Network Analysis of EOPE and LOPE

We analyzed 39 EOPE and 19 LOPE-associated exomiR networks to assess their systemic (Figure S1) and brain-specific impacts (Figure 1). These exomiRs were reported by Pillay *et al*. (Pillay et al., 2019). Systematically, network size varied widely: nodes ranged from 90 to 13,837, and edges ranged from 89 to 81,315, reflecting the heterogeneity and complexity of these interactions. The largest networks associated with has-miR-3613-3p in EOPE, contained 13,837 nodes and 81,315 edges, representing a highly connected systems linked to multiple interrelated molecular processes. In contrast, the smaller networks (i.e., has-miR151a-5p) consisted of 90 nodes and 89 edges, indicating more specific systems with a limited functional scope (Figure S1A). A similar pattern was observed in LOPE, where the most extensive network, hsa-miR-4516, included 10,989 nodes and 35,856 edges, while the most minor, hsa-miR-151b, contained 90 nodes and 89 edges (Figure S1B).

To evaluate the functional importance of brain-specific exomiR targets, we applied a composite metric (LogFC × mean expression in log₂ scale), which highlights proteins that are both strongly dysregulated and abundantly expressed, increasing their likelihood of biological impact. In the brain-specific networks (Figure 1B and 1C), EOPE interactomes contained between 80 and 10,140 nodes and 78 to 56,877 edges. In comparison, LOPE networks ranged from 80 to 8,161 nodes and 78 to 25,217 edges, indicating substantial structural variability. Despite this heterogeneity, high composite scores were observed in both small and large networks (1.92–16.30 EOPE; 2.84–6.93 LOPE), suggesting functionally relevant regulatory signals are maintained regardless of network size.

In the brain-specific analysis, EOPE networks exhibited a wide range of connectivity (0–1.67), indicating that their protein targets varied from weakly connected to highly integrated within the network. This modular and complex architecture likely reflects coordinated biological processes within tightly clustered protein groups. Within this framework, hsa-miR-3613-3p emerged as a central regulatory hub. By contrast, LOPE networks showed slightly higher overall density but greater structural heterogeneity (0–1.20), indicating that although proteins were, on average, more interconnected, their organization was uneven across modules. LOPE networks were predominantly structured around hsa-miR-4516, highlighting a distinct regulatory strategy compared with EOPE.

We next selected twelve exomiRs with the most significant differential expression and network complexity, defined by a high number of nodes and edges (Figure 1D). These included six EOPE-associated exomiRs (three upregulated and three downregulated; Figure 1B) and six LOPE-associated exomiRs (three upregulated and three downregulated; Figure 1C). All twelve selected exomiRs regulated proteins expressed in all 14 anatomical brain regions, highlighting widespread regulatory influence (Figure 1D).

### Functional and Subcellular Profiling of EOPE-and LOPE-Associated ExomiRs

Functional enrichment analysis of the six differentially expressed exomiRs in EOPE and LOPE (Figure 1D) revealed significant effects across biological processes, molecular functions, and cellular components when the whole human body proteome was considered (Figure S2 to S5).

In EOPE, upregulated exomiRs such as hsa-miR-223-3p (Figure S2) were enriched in pathways related to RNA metabolism, signaling, and transcriptional regulation. In contrast, downregulated exomiRs, including hsa-miR-3613-3p (Figure S3), were associated with stress response, ubiquitin–protein binding, and ribonucleoprotein complexes.

In LOPE, upregulated exomiRs such as hsa-miR-497-5p (Figure S4) and downregulated exomiRs such as hsa-miR-4516 (Figure S5) influenced pathways related to ubiquitin–protein ligase binding and focal adhesion. Across both EOPE and LOPE, exomiR-regulated proteins showed predominant subcellular localization in the cytosol, nucleus, and vesicles (>20%), consistent with roles in transcriptional and post-transcriptional regulation, vesicle trafficking, and oxidative stress (Figures S4–S5).

### Brain Homeostasis and ExomiR Dysregulation in EOPE

The six microRNAs differentially expressed in placental extracellular vesicles from women with EOPE were analyzed using the brain platform. Upregulated exomiRs were prioritized based on expression levels and bioinformatic relevance, which in the brain platform were related to neurovascular pathways, including inflammation, synaptic plasticity, oxidative stress, and BBB regulation. hsa-miR-223-3p (Figure 2A, Figure S6A and S6B) was associated with RNA metabolism and transcriptional regulation, with predicted localization to nuclear compartments. hsa-miR-16-5p (Figure 2B, Figure S6C and S6D) and hsa-miR-874-3p (Figure 2C, Figure S6E and S6F) were linked to intracellular transport and ubiquitin signaling, with enrichment in cytosolic, nuclear, and vesicular domains.

**Figure 2:**
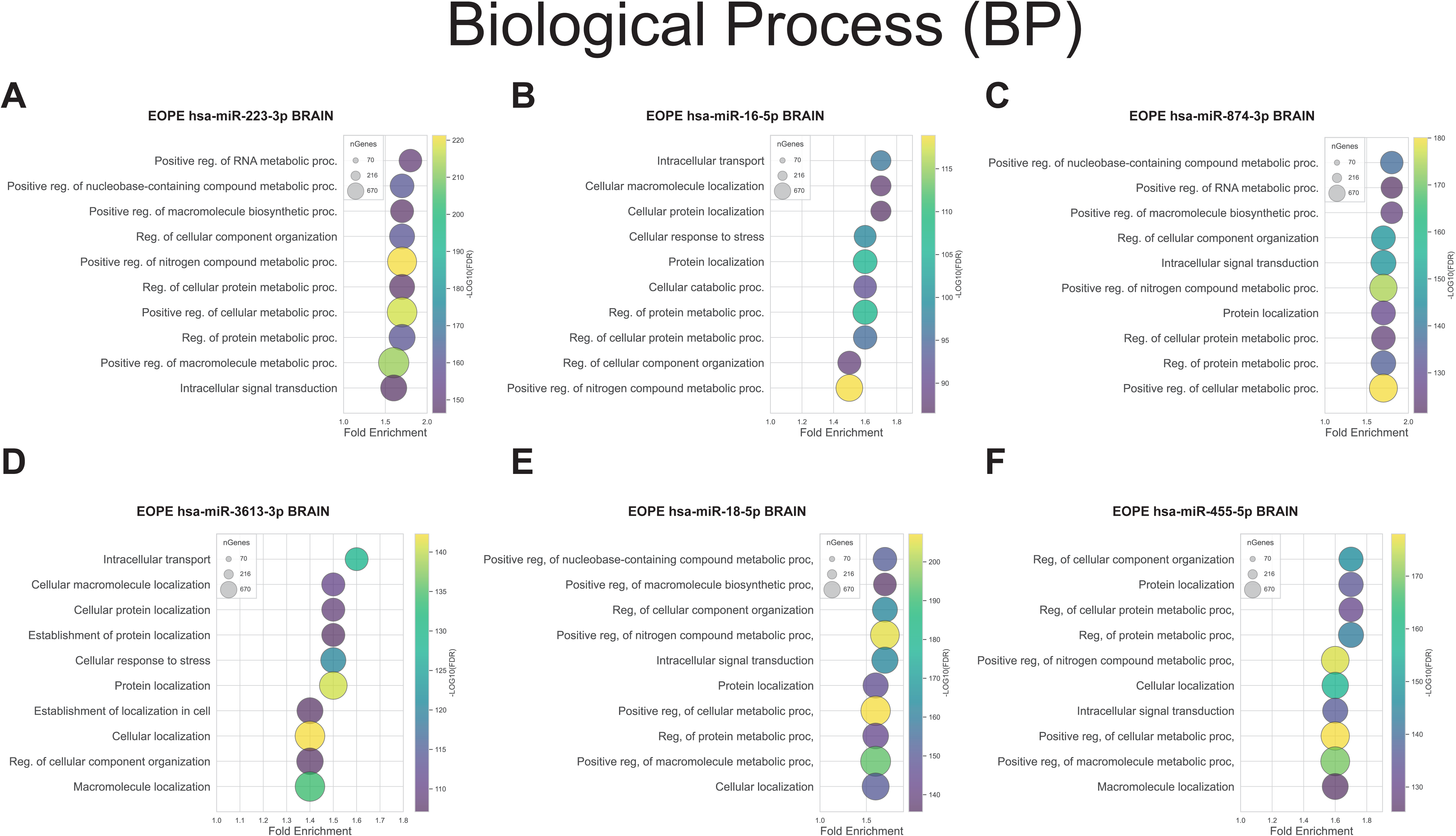
Functional Enrichment of Brain Proteins Targeted by EOPE-Associated ExomiRs. This plot illustrates the top 10 Gene Ontology (GO) terms with the most significant p-values, representing biological processes enriched by proteins modulated by early-onset preeclampsia (EOPE)-associated exomiRs. Each dot’s color indicates the p-value, and its size reflects the proportion of significantly differentially expressed genes (adjusted p-values <0.05) contributing to that specific GO term within each condition.

Downregulated exomiRs in EOPE showed overlapping roles in cellular stress response, biosynthesis, and structural organization. hsa-miR-3613-3p (Figure 2D, Figure S7A and S7B) was involved in nitrogen compound metabolism and organelle organization and localized to nuclear and ribonucleoprotein complexes. hsa-miR-18a-5p (Figure 2E, Figure S7C, and S7D) regulates RNA biosynthesis and kinase activity, and is enriched in chromatin and synaptic regions. hsa-miR-455-5p (Figure 2F, Figure S7E and S7F), which functionally resembles hsa-miR-18a-5p, exhibited distinct localization to focal adhesions, postsynaptic sites, and cell-substrate junctions, suggesting a specific role in neuronal connectivity and adhesion.

Collectively, these six exomiRs displayed heterogeneous subcellular distribution across 14 compartments, with predominant localization in the cytosol (18–20%), nucleus (12.5–17.5%), and vesicles (>20%). Their compartment-specific targets—hsa-miR-223-3p in the nucleus, hsa-miR-16-5p in mitochondria, hsa-miR-874-3p in synaptic vesicles, and hsa-miR-455-5p in postsynaptic domains— support a model of compartmentalized dysregulation in EOPE. Upregulated exomiRs may likely amplify inflammatory and oxidative stress signals, while downregulated exomiRs may impair transcriptional regulation, metabolic balance, and endothelial adaptation, contributing to neurovascular dysfunction in EOPE.

### Brain Homeostasis and ExomiR Dysregulation in LOPE

The six exomiRs differentially expressed in LOPE, namely hsa-miR-497-5p, hsa-miR-32-5p, hsa-miR-202-3p, hsa-miR-4516, hsa-miR-488-3p, and hsa-miR-494-5p, were further analyzed and found to regulate critical pathways involved in oxidative homeostasis, mitochondrial function, and intracellular trafficking (Figure 3). Functional enrichment analysis identified roles in nitrogen metabolism, vesicle transport, and structural organization, with gene ontology terms reflecting regulation of cellular components and responses to stress. These exomiRs are involved in processes related to synaptic maintenance, proteostasis, and BBB integrity.

**Figure 3:**
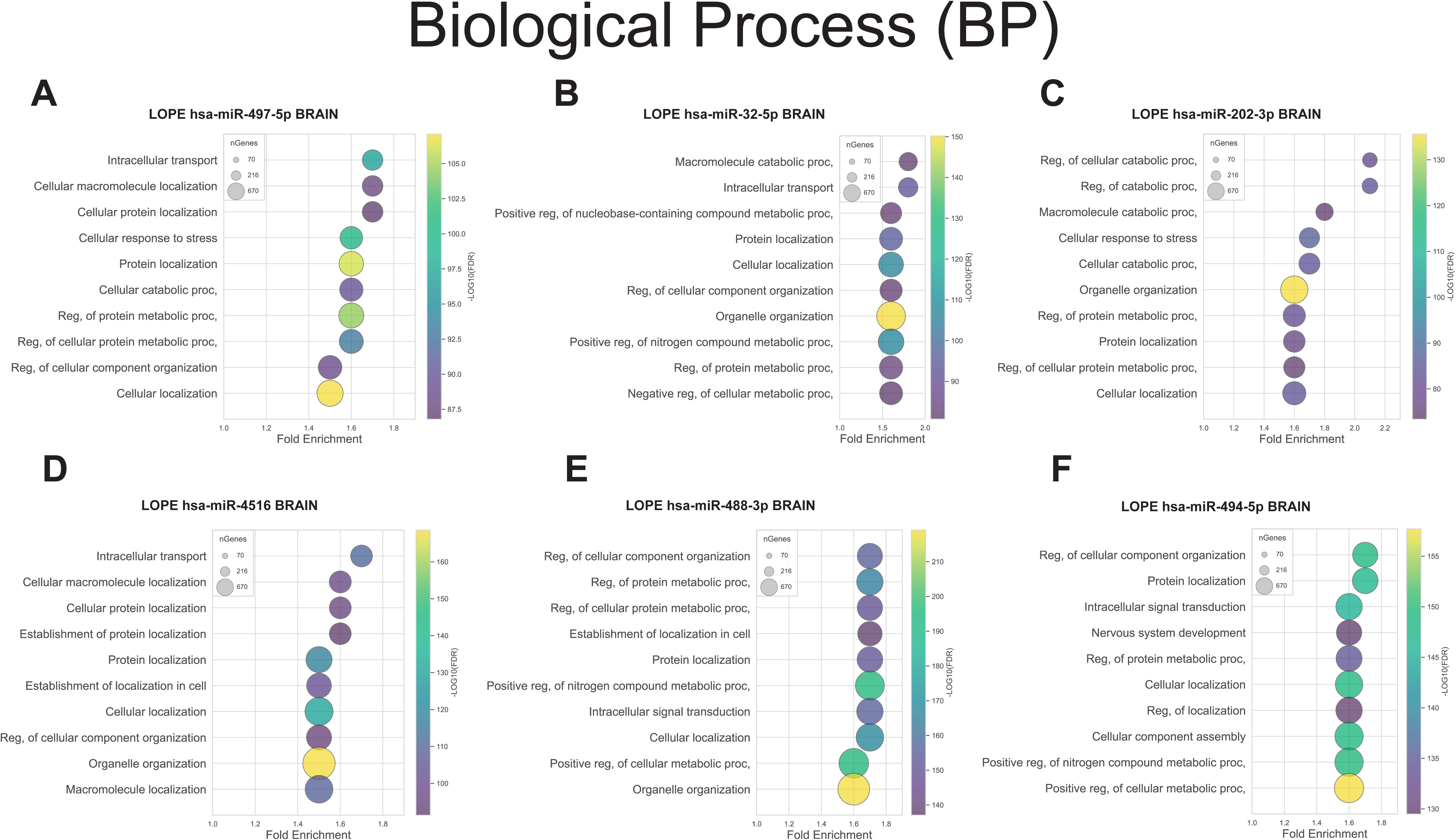
Functional Enrichment of Brain Proteins Targeted by LOPE-Associated ExomiRs. This figure displays the top 10 Gene Ontology (GO) terms with the highest statistical significance (lowest p-values), indicating biological processes enriched among proteins regulated by late-onset preeclampsia (LOPE)-associated exomiRs. Dot color corresponds to the p-value, and dot size is scaled to the count of significantly differentially expressed genes (adjusted p-values <0.05) mapping to each GO term across conditions. All data utilized were derived from biological process annotations.

For example, hsa-miR-202-3p (Figure 3C, Figure S8E and S8F) and hsa-miR-494-5p (Figure 3F, Figure S9E and S9F) may influence synaptic and endothelial functions, while hsa-miR-4516 (Figure 3D, Figure S9A and S9B) may regulate proteostasis and BBB stability (Table S1). Subcellular localization analysis revealed that these exomiRs are predominantly localized to the cytosol (18-20%), nucleus (13-18%), and vesicles (>20%), consistent with their involvement in transcriptional and post-transcriptional regulation, vesicle trafficking, and responses to oxidative stress. hsa-miR-4516 localized to mitochondria, while hsa-miR-494-5p and hsa-miR-488-3p were enriched in cytoskeletal and membrane compartments.

Taken together, both EOPE and LOPE-associated exomiRs converge on pathways involving mitochondrial dysfunction and cytoskeletal instability—hallmarks also implicated in neurodegenerative disease. Key functional clusters of exomiRs and their associated pathways are summarized in Table S1.

### Proteomic Signatures in Cerebrospinal Fluid (CSF)

To explore the link between placental exomiRs and maternal brain alterations, we integrated the six differentially expressed EOPE exomiRs with CSF proteomic data (Ciampa et al., 2018). In silico analyses focused on first-order exomiR targets that overlapped with proteins differentially expressed in the CSF of women with EOPE, either compared with healthy controls or with EOPE patients presenting neurological complications (Figure 4A).

**Figure 4.**
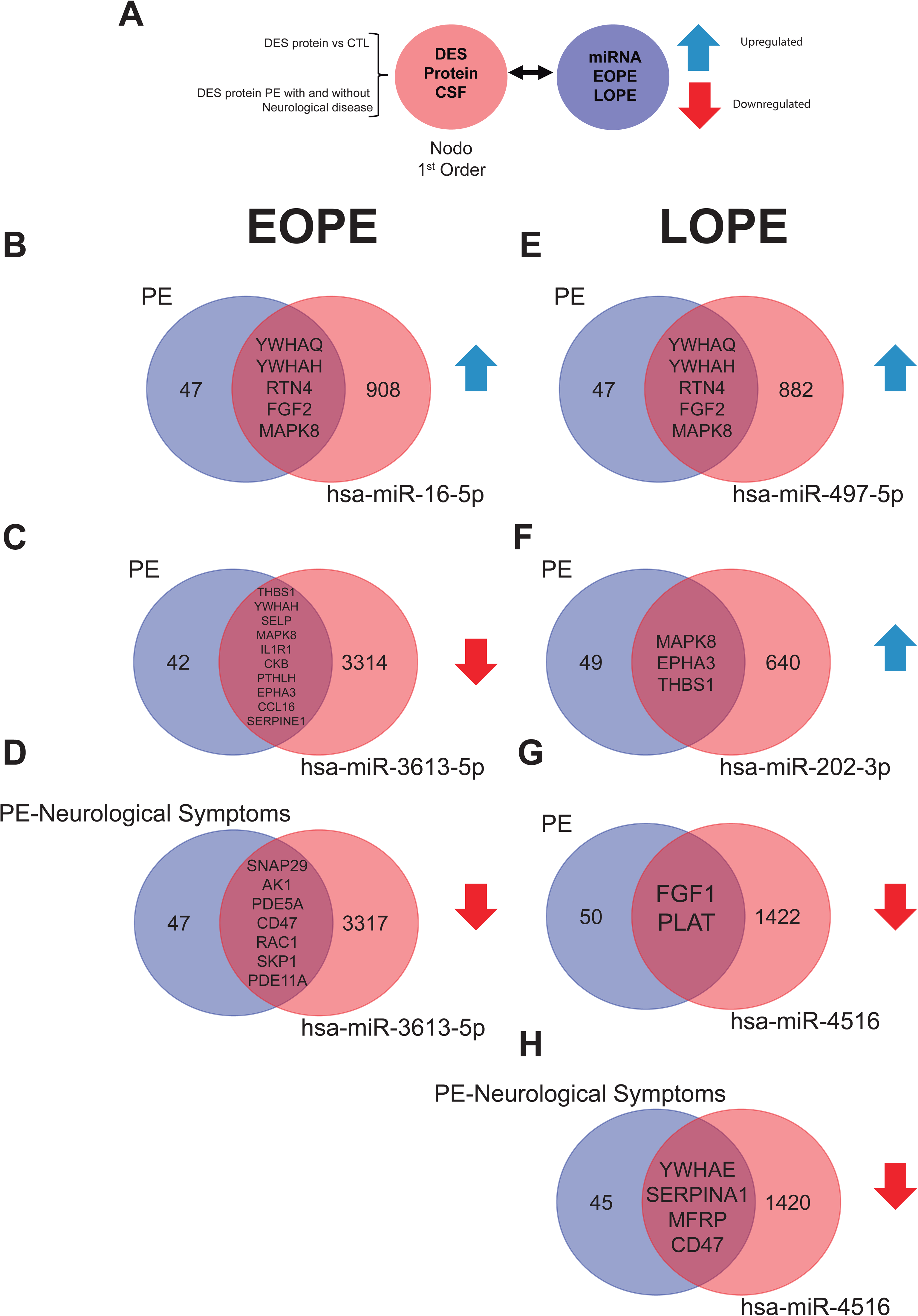
Cerebrospinal fluid (CSF) proteomic convergence of preeclampsia (PE)-associated ExomiRs. (A) Workflow of exomiR–protein interaction mapping. (B–H) Networks of differentially expressed proteins intersecting with selected exomiRs (e.g., hsa-miR-16-5p, hsa-miR-3613-5p, hsa-miR-497-5p, hsa-miR-202-3p, hsa-miR-4516). Key nodes include YWHA family members, MAPK8, FGF2, and THBS1, highlighting pathways associated with neurodegeneration and cerebrovascular instability in preeclampsia. DES, differentially expressed proteins. CTL, control. Preeclampsia, PE.

Among the 6 chosen differentially expressed miRNAs in EOPE, only two (has-miR-16-5p, and has-miR-3613-5p) (Figure 4B and 4C) converge with the first node interactions with differentially expressed proteins in the CSF of EOPE with respect to control. miR-3613-5p exhibited twice the interactions with 10 overlapping protein targets compared with miR-16-5p. From them, common proteins included MAPK8 and YWHAH.

When analysis was restricted to EOPE exomiRs in PE patients who had neurological symptoms, only has-miR-3613-5p (Figure 4D) overlapped with CSF proteins targeting seven unique proteins: SNAP29, AK1, PDE5A, CD47, RAC1, SKP1, and PDE11A. Although diverse in their individual functions, these proteins collectively contribute to critical brain functions, including synaptic transmission, signal modulation, energy metabolism, and immune regulation. SNAP29 and RAC1 are key players in vesicle trafficking and cytoskeletal dynamics, while PDE5A and PDE11A modulate intracellular signaling through cyclic nucleotides, influencing processes like synaptic plasticity. AK1 supports neuronal energy metabolism, and CD47 regulates immune signaling, impacting neuroinflammation and synaptic pruning. Their coordinated roles underline the complexity of neuronal maintenance and brain health, with potential implications for neurodegenerative diseases and cognitive dysfunctions.

In LOPE, three exomiRs—hsa-miR-497-5p (Figure 4E), hsa-miR-202-3p (Figure 4F), and hsa-miR-4516 (Figure 4G)—interacted with differentially expressed CSF proteins compared with controls. These exomiRs shared five, three, and two targets, respectively, with MAPK8 again emerging as a standard hub for both hsa-miR-497-5p and hsa-miR-202-3p. Of note, both belonged to the upregulated LOPE exomiR group.

Significantly, hsa-miR-4516 interacted with differentially expressed proteins in CSF of women with LOPE who also exhibited neurological symptoms (Figure 4H). Common target proteins included YWHAE, SERPINA1, MFRP, and CD47. Those proteins may contribute to brain function through roles in signaling, neuroprotection, development, and immune regulation. For instance, YWHAE modulates synaptic plasticity, SERPINA1 provides anti-inflammatory and neuroprotective effects, MFRP supports neuronal development via Wnt signaling, and CD47 regulates neuroimmune interactions.

Overall, integrating EOPE and LOPE exomiRs identifies five placental exomiRs (of the twelve that were initially chosen, Figure 1D) that share brain differentially expressed targets detected in CSF proteomic profiles. These exomiRs regulate proteins involved in neuroprotection (YWHA family proteins, FGF2), synaptic plasticity (THBS1, EPHA3), and endothelial signaling (SELP, IL1R1), suggesting their contribution to inflammation, vascular instability, and neuronal dysfunction. Moreover, recurrent targets such as MAPK8 and FGF2 across multiple exomiRs point to shared molecular mechanisms that may contribute to cerebrovascular and cognitive impairments in PE, particularly in late-onset presentations (Table S2).

### Regulation of Blood-Brain Barrier Integrity

Claudin 5 (CLDN5) is a critical tight junction protein that maintains BBB integrity (Angelow et al., 2008), with high expression in brain endothelial cells (Greene et al., 2019; Morita et al., 1999). Preclinical studies have shown that extracellular vesicles derived from hypoxic placental cultures reduce CLDN5 expression at the BBB (Sandoval et al., 2025). Based on this, we analyzed interactions between exomiRs and a curated 43-protein CLDN5 network (Figure 5A). These interactions converge on mechanisms that affect tight junction stability, endothelial signaling, and vesicle transport—key factors in BBB disruption in PE (Table S3).

**Figure 5.**
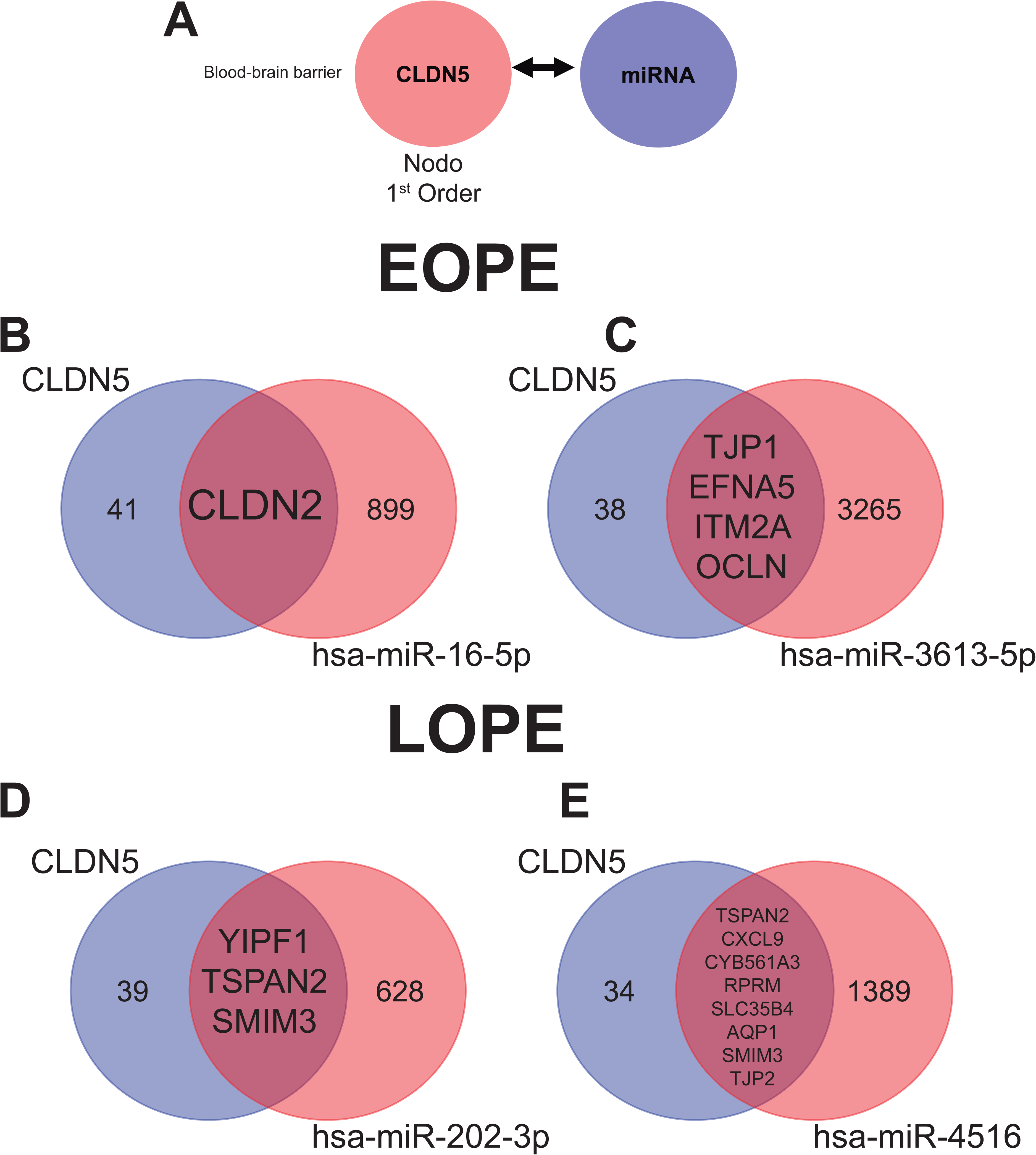
ExomiR interactions with the CLDN5 tight-junction network at the blood–brain barrier. (A) Schematic of the curated first–order CLDN5 network (43 proteins) used to test overlaps with PE-associated exomiRs. (B–C) EOPE-associated exomiRs (hsa-miR-16-5p, hsa-miR-3613-5p) share key proteins, such as CLDN2, TJP1, and OCLN, suggesting mechanisms of tight-junction destabilization. (D–E) LOPE-associated exomiRs (hsa–miR–202–3p, hsa–miR–4516) overlap with multiple proteins (e.g., TSPAN2, AQP1, TJP2), indicating broader regulation of endothelial integrity. Numbers in the Venn diagrams denote unique or shared proteins. DES, differentially expressed proteins. CTL, control. Preeclampsia, PE.

Four exomiRs (hsa-miR-16-5p, hsa-miR-3613-5p, hsa-miR-202-3p, and hsa-miR-4516) targeted proteins essential for BBB structure and regulation (Figure 5B–5E). For example, hsa-miR-16-5p interacted with CLDN2 in proximity to CLDN5, while hsa-miR-3613-5p regulated TJP1, EFNA5, ITM2A, and OCLN in EOPE (Figure 5C). In LOPE, hsa-miR-4516 displayed the broadest impact, targeting eight CLDN5-associated proteins (Figure 5E), whereas hsa-miR-202-3p affected only three (Figure 5D). Common targets across EOPE and LOPE included TSPAN2 and SMIM3, both of which are implicated in vesicle trafficking, membrane stability, and junctional disassembly.

Together, these findings suggest that EOPE exomiRs destabilize the BBB through coordinated regulation of multiple tight junction proteins. In contrast, LOPE exomiRs, dominated by hsa-miR-4516, act through a more centralized hub-driven mechanism. These distinct strategies point to different molecular pathways of neurovascular injury in PE.

### Neuroanatomical Impact of ExomiRs: Regional and Cell-Type-Specific Protein Regulation in Preeclampsia

Figure 6 summarizes how exomiRs associated with PE may modulate brain protein expression across distinct regions and cell types, offering insight into the neurological manifestations of the condition.

**Figure 6.**
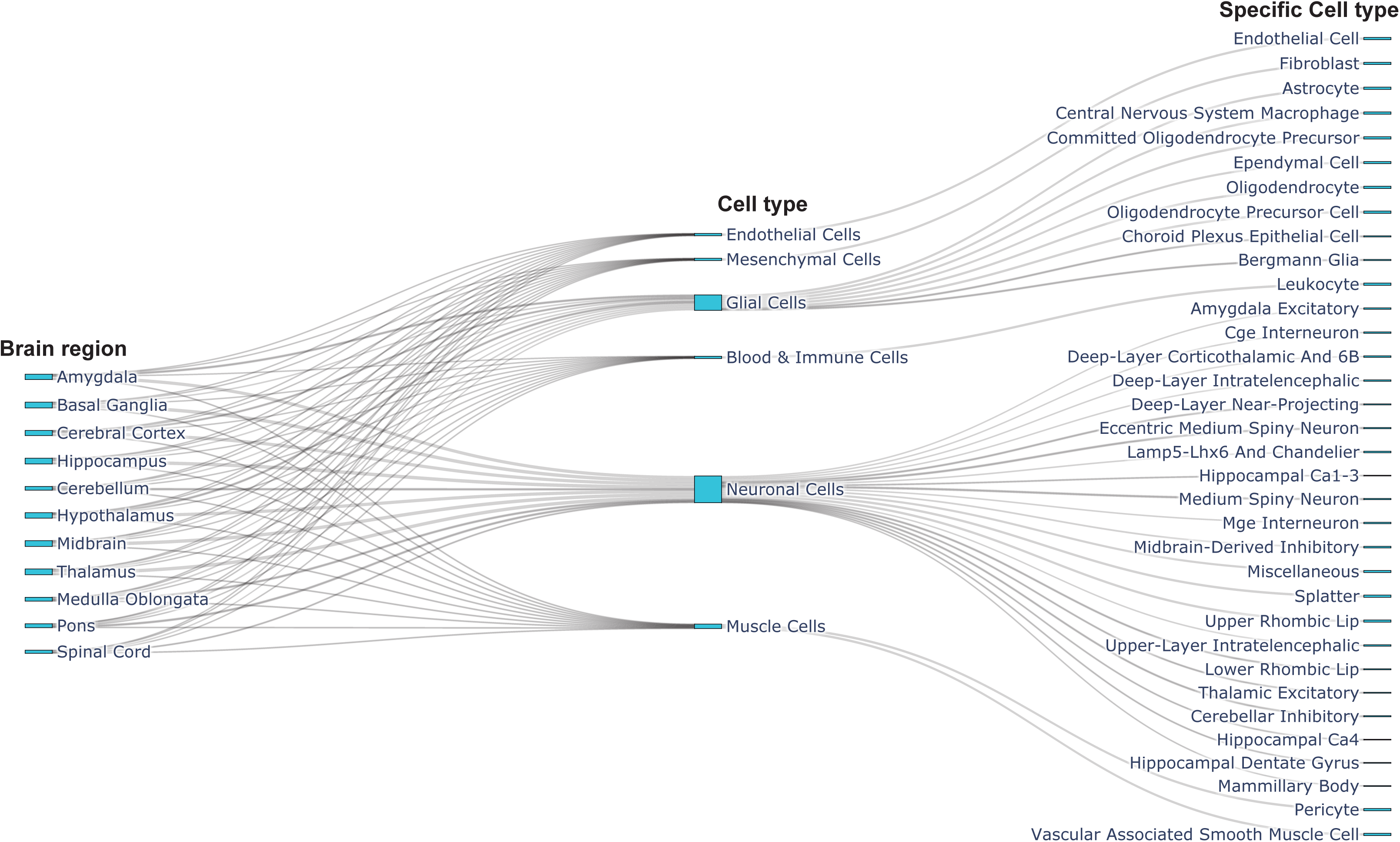
Neuroanatomical mapping of exomiR-regulated proteins in preeclampsia. ExomiRs modulate proteins across the amygdala, cerebellum, cortex, hippocampus, and motor regions, affecting endothelial cells, astrocytes, interneurons, and microglia, disrupting the blood-brain barrier (BBB), synaptic plasticity, and neuromuscular function.

In the amygdala, exomiR-targeted proteins in endothelial cells, fibroblasts, astrocytes, and central nervous system (CNS) macrophages suggest compromised BBB integrity, extracellular matrix disorganization, glial stress, and neuroinflammation mechanisms likely contributing to emotional and cognitive disturbances. In the cerebellar gyrus, altered expression of GABAergic interneuron enzymes may impair inhibitory signaling and cerebellar coordination. In the frontal cortex, dysregulation in pyramidal neurons and astroglia may affect synaptic plasticity, glial activation, and vascular homeostasis. ExomiR targets in hippocampal granule neurons involve key proteins for memory and glutamatergic signaling, linking PE to cognitive dysfunction. In the motor cortex, altered regulation of acetylcholine synthesis and calcium channel proteins suggests impaired neuromuscular function.

Together, these findings reveal a spatially resolved pattern of exomiR-mediated protein dysregulation impacting inflammation, neurovascular and synaptic function, and metabolic stability. This neuroanatomical framework highlights exomiRs as key contributors to PE-related brain injury, providing mechanistic targets for biomarker and therapeutic development.

## Discussion

Cerebrovascular complications remain one of the leading causes of maternal death in women with PE, and survivors face an elevated risk of long-term neurocognitive decline. Our study expands current understanding by demonstrating how placental exosomes released into the maternal circulation may target the maternal brain, underscoring the vast potential for placenta–brain interactions. By integrating multi-level network analysis with CSF proteomics, we dissected brain-specific regulatory effects of PE-associated exomiRs, providing novel insights into their potential roles in neurovascular dysfunction and cognitive risk.

Emerging evidence increasingly implicates placental-derived exomiRs as a potential pivotal mediator of systemic and neurovascular dysfunction in PE, extending beyond the traditional boundaries of obstetric pathology. Building on the foundational work by Pillay et al. (Pillay et al., 2019) and Awoyemi et al. (Awoyemi et al., 2024), our study demonstrates that exomiRs derived from circulating EOPE and LOPE exhibit distinct molecular signatures with profound implications for maternal brain health. These signatures not only predict placental dysfunction but also appear to orchestrate long-term neurovascular and neurodegenerative consequences, directly supporting the concept of a placenta-brain axis.

Consistent with Pillay *et al*. (Pillay et al., 2019), we confirmed that EOPE and LOPE are molecularly distinct entities, characterized by heterogeneous exomiR regulatory landscapes. EOPE exomiRs were enriched in pathways related to lipid metabolism, inflammation, and thrombosis, displaying broader connectivity and higher node complexity. Conversely, LOPE exomiRs were associated with metabolic dysregulation and cellular stress responses. These findings align with the pathophysiological characterization of EOPE as a more severe disorder of shallow placentation and placental ischemia, and LOPE as a disease of placental senescence and oxidative stress (Awoyemi et al., 2024; Pillay et al., 2019). Importantly, our brain-specific network analysis revealed that these exomiRs (at least the twelve selected in this analysis) regulate proteins involved in BBB integrity, synaptic plasticity, and neuroinflammation, suggesting a direct mechanistic link between placental dysfunction and maternal brain injury.

Our analyses focused on twelve highly differentially expressed exomiRs, six from EOPE (hsa-miR-223-3p, hsa-miR-16-5p, hsa-miR-874-3p, hsa-miR-3613-5p, hsa-miR-18a-5p, hsa-miR-455-5p) and six from LOPE (hsa-miR-497-5p, hsa-miR-32-5p, hsa-miR-202-3p, hsa-miR-4516, hsa-miR-488-3p, hsa-miR-494-5p), all these exomiRs were previously reported by Pillay *et al*. (Pillay et al., 2019). This prioritization enables in-depth analysis of brain-specific protein networks across all 14 anatomical brain regions. These exomiRs targeted proteins involved in tight junction dynamics (e.g., CLDN5, TJP1), synaptic transmission (e.g., SNAP29, YWHAH), and neuroinflammation (e.g., CD47, RAC1). Notably, hsa-miR-3613-5p and hsa-miR-4516 emerged as central hubs in both EOPE and LOPE networks, suggesting their potential as critical biomarkers and therapeutic targets. For example, EOPE-upregulated exomiRs (e.g., hsa-miR-223-3p and hsa-miR-16-5p) were linked to nuclear and mitochondrial regulatory processes, potentially amplifying oxidative stress and impairing transcriptional homeostasis. Conversely, downregulated EOPE exomiRs (e.g., hsa-miR-3613-3p) appear to disrupt biosynthetic and stress response pathways, collectively compromising neurovascular integrity.

Our integration with CSF proteomics further validated these findings and reinforced the existence of a robust placenta–brain axis. ExomiRs from both EOPE and LOPE placentas converged on shared brain protein targets, including MAPK8, FGF2, and YWHA family proteins, all of which are implicated in neuroprotection, synaptic plasticity, and vascular stability. The recurrent targeting of MAPK8 by multiple exomiRs underscores its role as a nodal regulator of neuroinflammation and oxidative stress, a mechanism previously described in PE-associated brain endothelial dysfunction (Cipolla et al., 2012; Johnson et al., 2014; Li et al., 2017). In addition, hsa-miR-3613-5p displayed strong interactions with proteins involved in vesicle trafficking, cytoskeletal dynamics (e.g., SNAP29, RAC1), and immune modulation, which could exacerbate BBB dysfunction, a hallmark of PE-associated neurological complications (Friis et al., 2022; Johnson et al., 2014; Warrington et al., 2015).

Neuroanatomical mapping of exomiR targets revealed region-and cell-type-specific vulnerabilities. In the amygdala and hippocampus, exomiRs targeted proteins in endothelial cells and astrocytes, suggesting that BBB disruption and glial stress are key mechanisms. Conversely, cerebellar and cortical regions showed enrichment for proteins involved in synaptic signaling and neuromuscular control, aligning with clinical observations of cognitive deficits in PE survivors (Aukes et al., 2008; Beckett et al., 2023; Carey et al., 2024). These findings corroborate recent studies linking placental exomiRs to neurodegenerative phenotypes, including increased β-amyloid and tau pathology in postpartum women with prior PE (Suvakov et al., 2023).

Mechanistically, the pathways leading to BBB impairment in PE remain incompletely understood. Several circulating factors have been linked with preclinical evidence of BBB disruption, including VEGF, TNF-α, LDL oxidized, or Angiotensin II type 1-Receptor Autoantibodies (AT1-AA) (Bergman et al., 2021; Duncan et al., 2020; Johnson et al., 2018; Schreurs and Cipolla, 2014), as well as placental EVs (Leon et al., 2021; Sandoval et al., 2025). We recently demonstrated that EVs isolated from a placenta cultured under hypoxia, as a proxy of PE, decrease CLDN5 functional expression in brain endothelial cells (Sandoval et al., 2025). Notably, unlike BBB disruption induced by plasma from women with PE (Bergman et al., 2021; Johnson et al., 2018), the EV-mediated effect on CLDN5 could not be reversed with inhibition of VEGF receptor 2 (Sandoval et al., 2025). These findings suggest that other EV cargo, particularly exomiRs, may contribute to BBB disruption by downregulating the functional expression of CLDN5.

Our study, therefore, bridges the gap between obstetric pathology and neurology by demonstrating that placental exomiRs are not merely by-products of PE but may act as active mediators of neurovascular injury. Furthermore, the compartmentalized dysregulation of exomiRs, particularly their enrichment in synaptic vesicles, mitochondria, and nuclear compartments, supports a model of spatially constrained neurotoxicity, where exomiRs may exert cell-type and region-specific effects.

In conclusion, our study provides bioinformatic evidence that placental-derived exomiRs are instrumental in the pathophysiology of cerebrovascular complications associated with PE. Our work delineated a robust molecular framework in which exomiRs act as systemic and neurovascular effectors, mediating BBB disruption, synaptic dysfunction, and neuroinflammation. The identification of shared and distinct exomiR signatures in EOPE and LOPE not only reinforces the heterogeneity of PE but also offers a precise roadmap for diagnostics and therapeutic interventions. Our findings highlight the dual role of exomiRs as both promising biomarkers and active mediators of neurovascular pathology, firmly supporting the concept of a placenta-brain axis in PE. Future studies should validate these candidate exomiRs in larger longitudinal cohorts and explore their utility as early diagnostic tools for neurodegenerative risks associated with PE, as well as elucidate their long-term impact on cognitive and vascular health.

## Acknowledgment

The authors would like to thank the researchers from GRIVAS Health and RIVATREM collaborative networks.

## Author contributions

MA conceptualized the study. CE led the research team. MV were consultants in exosome/sEVs characterization and data analysis. MA, CE, and CA edited the manuscript. All co-authors approved the final version of this manuscript.

## Supplemental Material

Expanded Materials and Methods

Figures 1-9

